# Phenotypic plasticity segregates antagonistic species but undermines their coexistence

**DOI:** 10.64898/2026.11.03.763922

**Authors:** Diya Elizabeth Shaji, Muhammed Hadi, Yashraj Chavhan

## Abstract

Environmental change often induces novel phenotypes in multiple cohabiting species, yet how such common-cue-induced plasticity shapes their coexistence is largely unknown. Testing this link requires disentangling inducing-environment effects from induced-phenotype effects, which remains elusive. We address this gap using two mutually antagonistic bacterial species, *Staphylococcus aureus* (*Sa*) and *Pseudomonas aeruginosa* (*Pa*). In habitual salinity, where both bacteria show unicellularity, *Pa* severely diminished *Sa*. In contrast, high salinity induced macroscopic cell clustering that spatially segregated *Sa* and *Pa* (*Sa*: bottom-dwelling vertical filament; *Pa*: top-dwelling horizontal mat), and led to >10-fold higher coexistence than habitual salinity. Counterintuitively, physically inhibiting cell clustering (abolishing segregation) in high salinity led to a further >100-fold higher coexistence (*Sa*:*Pa*∷∼1:1). This was because salinity-induced clustering was adaptive for *Pa*, but not for *Sa*: such adaptive plasticity enabled killer *Pa* to proliferate by monopolizing oxygen, which diminished *Sa* in a correlated manner, thereby weakening coexistence (mechanistically explained by diffusion-based killing). Hence, although the inducing environment promoted overall coexistence, the induced phenotype undermined it. Community-wide genomics revealed that the high level of coexistence achieved by abolishing segregation in high salinity allowed rapid coculture-specific evolution. Taken together, we uncover how plasticity can appear to promote antagonistic coexistence despite causally undermining it.

## Introduction

Phenotypic plasticity, the capacity of a single genotype to exhibit different phenotypes in contrasting environments, is a ubiquitous feature of the living world^1–4^. In principle, such plasticity can be adaptive or maladaptive^5–9^, bringing organisms closer to or farther from fitness optima in altered environments. Empirical studies linking plasticity to fitness effects largely rely on detecting the *co-occurrence* of two changes: (1) a changed phenotype induced by the new environment and (2) altered fitness in this new environment^10–12^. However, this approach faces a key conceptual hurdle: If an environmental change leads to both a new phenotype and an altered fitness, this does not imply that the new phenotype *causes* the fitness change. Such altered fitness may arise directly from the environment itself rather than from the induced phenotype. Moreover, most theoretical and empirical studies link plastic phenotypes to fitness changes in a single species^8,10,11,13^. But species are rarely found in isolation, and much less is known about the fitness effects of plasticity induced simultaneously in multiple interacting species by a common environmental cue within an ecological community^11^. This gap is particularly evident for experimental studies of plasticity in mutually antagonistic species^11,14^, especially when the plastic phenotypes transcend multiple generations and can have asymmetric fitness consequences across distinct species^15–18^. Beyond such asymmetric effects, plasticity can significantly alter interspecies interactions, which can cascade through the community^11,19^. Given these multivariate challenges, predicting something as fundamental as whether plasticity induced by a common cue improves or hinders interspecies coexistence becomes very difficult, even in a simple community with only two antagonistic species^11,20,21^.

In principle, plasticity can have both positive and negative effects on antagonistic coexistence^11,20–23^, depending on how it shapes two key aspects across species: niche contrasts and fitness asymmetries. Most intuitively, plasticity can promote coexistence by segregating antagonistic species in space and/or time, thereby reducing their overlap (and hence, interactions)^20,24–27^. Analogously, if plasticity increases the spatiotemporal overlap of antagonistic species in terms of their ecological niches, it can lead to reduced antagonistic coexistence^22,11,28^. Furthermore, in a given pair of antagonistic species, if plasticity benefits one species much more than the other, it has the potential to reduce coexistence^29^. Although multiple empirical studies have documented such links between plasticity and antagonistic coexistence, their inferences rely on the co-occurrence of phenotypic and fitness changes in altered environments^11^. Indeed, a recent important study connecting plasticity to interspecies coexistence highlighted that a key empirical challenge in this area is to incorporate a treatment expressing plasticity versus another not expressing it^20^. Importantly, the key causal uncertainty identified above has not been tackled directly: whether a change in antagonistic coexistence is driven directly by plasticity, or more parsimoniously, whether such change is caused by the altered environment itself. Here we directly addresses this causal ambiguity using a pair of mutually antagonistic bacteria, *Staphylococcus aureus* (henceforth, *Sa*) and *Pseudomonas aeruginosa* (henceforth, *Pa*). These species routinely coexist in their natural habitat (chronic wounds and cystic fibrosis sputa in humans)^30–32^. However, they are extremely difficult to coculture in the laboratory because *Pa* tends to drive *Sa* to extinction in environments that are otherwise highly conducive for both species in isolation^31–35^.

Here we show that high environmental salinity induces macroscopic clustering in both *Sa* and *Pa*, which causes the spatial segregation of these antagonistic species within liquid cocultures. We also find that high salinity substantially improves the coexistence of these antagonistic bacteria. Although this co-occurrence of spatial segregation and improved *Sa*-*Pa* coexistence may suggest that spatial segregation (which is expected to reduce antagonistic interactions) *causes* the improved coexistence, we empirically show that such ecological intuition can be highly misleading. Using a series of experiments, we disentangle the role of the inducing environment from that of the induced phenotype, and demonstrate that these roles can oppose each other. We show that inhibiting plastic cell clustering (and thus abolishing spatial segregation) in high salinity leads to the highest level of coexistence where both the antagonistic species maintain fairly equal abundance over long periods and undergo coculture-specific molecular evolution. We further show that the causal basis of this surprising phenomenon is the uneven adaptive utility of phenotypic plasticity across the two species: High-salinity-induced cell clustering is adaptive for *Pa*, but not for *Sa*. We demonstrate that such adaptively plastic cell clustering allows *Pa* to proliferate by monopolizing oxygen, which results in a concomitant depletion in *Sa* counts in a negatively correlated manner that we mechanistically explain using diffusion-based killing experiments. This shows how high salinity (the inducing environment) increases the overall coexistence, but the induced phenotype (cell clustering) undermines it.

## Results

### High salinity spatially segregates *S. aureus* and *P. aeruginosa* in cocultures by inducing macroscopic cell clustering

We have recently shown that both *S. aureus* (*Sa*) and *P. aeruginosa* (*Pa*) exhibit phenotypically plastic macroscopic cell clustering^36^. Specifically, when grown as single-species monocultures in Luria Bertani (LB) broth with 0.5% NaCl (w/vol), both bacteria show unicellular planktonic growth. However, in LB with 6% NaCl (w/vol), both *Sa* and *Pa* monocultures grow as macroscopic clusters containing several million viable cells^36^. Following Ref. 36, we call these environments “habitual salinity” and “high salinity”, respectively. We first tested if such phenotypically plastic macroscopic clustering made *Pa* and *Sa* monocultures occupy similar physical niches within unshaken liquid cultures. To this end, we grew *Pa* and *Sa* monocultures in both habitual and high salinity for 3 days (see *Methods*). We found that in high salinity, *Sa* and *Pa* monocultures formed markedly different macroscopic clusters, differing in both the morphology and the spatial location (Fig. S1). Specifically, high salinity made *Sa* form an elongated vertical stalk emanating from the bottom of the culture tube (Fig. S1). In contrast, high salinity made *Pa* grow as a thick horizontal mat at the air-liquid interface (Fig. S1). As expected, neither *Sa* nor *Pa* showed noticeable macroscopic clustering under habitual salinity (Fig. S1). Building on these observations from *Sa* and *Pa* monocultures, we determined if *Sa* and *Pa* exhibited key contrasts in both the morphology and the location of their macroscopic clusters when they are grown together in cocultures. To this end, we grew unshaken cocultures with a 1:1 starting inoculum of *Sa* and *Pa* in both habitual and high salinity for 3 days (see *Methods*). The habitual salinity cocultures did not show noticeable cell clustering, as expected (Fig. 1a). Interestingly, the high salinity cocultures showed both an interface-inhabiting horizontal mat at the top and a vertical stalk emanating from the bottom (Fig. 1a). Next, we determined the composition of the mats and stalks from such high salinity cocultures (Fig. 1b; see *Methods*). In all the replicates, the horizontal mats were constructed almost exclusively by *Pa* (Fig. 1b). In stark contrast, the high salinity-induced vertical stalks in the cocultures were made almost exclusively by *Sa* (Fig. 1b). Thus, phenotypically plastic cell clustering spatially segregates *Sa* and *Pa* at macroscopic scales (Fig. 1b).

**Fig. 1.**
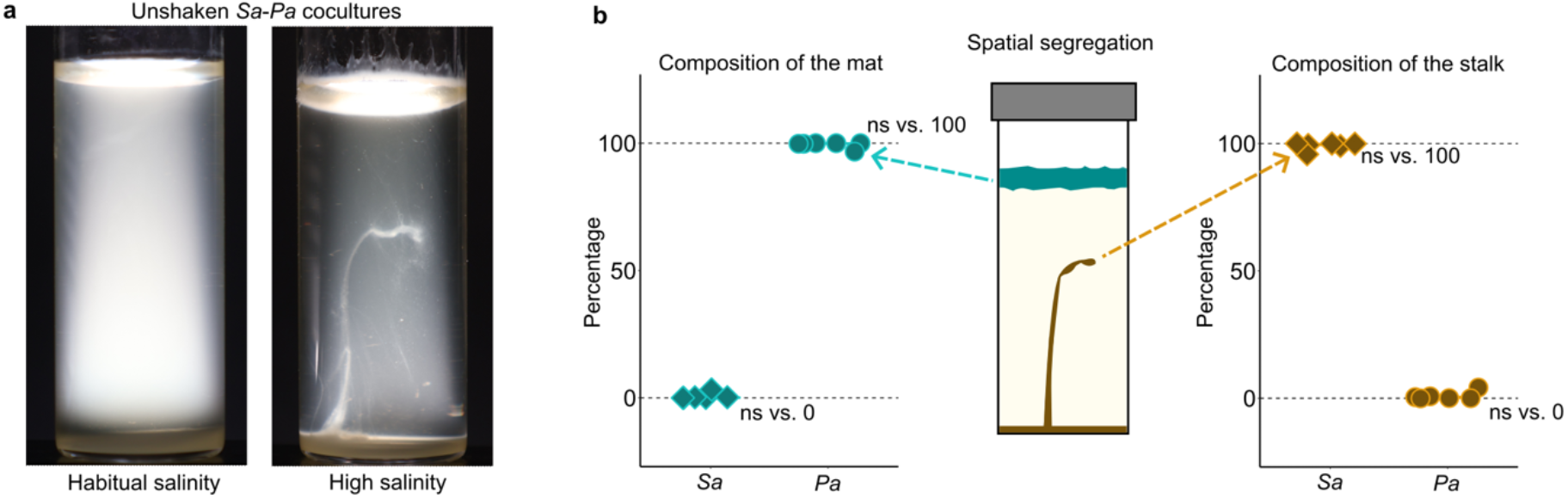
High salinity causes the macroscopic spatial segregation of *S. aureus* and *P. aeruginosa*. (a) The growth of *Sa*-*Pa* cocultures under habitual and high salinity environments. Whereas the habitual salinity cocultures were planktonic and turbid, the high salinity cocultures exhibited a horizontal mat at the air-liquid interface that was spatially distinct from a vertical stalk emanating from the bottom of the tube. (b) High salinity segregates *Pa* and *Sa* in space (see SA.1 in *Statistical Appendix* for details).

### High salinity promotes the coexistence of *S. aureus* and *P. aeruginosa*

*Pa* is known to drive *Sa* to extinction via interference competition mediated by active killing in conditions similar to our habitual salinity environment^31–35^. Thus, previously established literature suggests that under habitual salinity, a coculture initiated with a 1:1 *Sa*-*Pa* inoculum will become dominated by *Pa* while *Sa* will approach extinction. An environment that can reduce interactions between these antagonistic species has the potential to improve their coexistence. One of the simplest ways of reducing antagonistic interactions between two species is to segregate them spatially^37,22,38,39^. Based on this rationale, we hypothesized that because high salinity causes the spatial segregation of *Sa* and *Pa* (Fig. 1), high salinity will promote the coexistence of these antagonistic species. To test this hypothesis, we grew unshaken cocultures with a 1:1 starting inoculum of *Sa* and *Pa* under both habitual and high salinity for 7 days (see *Methods*). At the end of this 7-day period, we vortexed the culture tubes and determined the relative fraction of *Sa* and *Pa* within each community (see *Methods*). We found that in habitual salinity cocultures, the fraction of *Pa* was overwhelmingly larger than that of *Sa*, as expected (Fig. 2a). Aligning with our key prediction, high salinity led to a substantial increase in the coexistence of *Sa* and *Pa.* Specifically, the relative abundance of *Sa* was >10-fold higher in high salinity as compared to that in habitual salinity (Fig. 2a). Our usage of the relative abundance of *Sa* as a measure of coexistence is motivated by previous studies in microbial ecology^40,41^ (see *Methods*). Ecological intuition suggests that high salinity improves *Sa-Pa* coexistence *because* it spatially segregates *Sa* and *Pa* (Fig. 2b). We tested this intuitive notion by conducting a series of experiments in both high and habitual salinity (Fig. 3a).

**Fig. 2.**
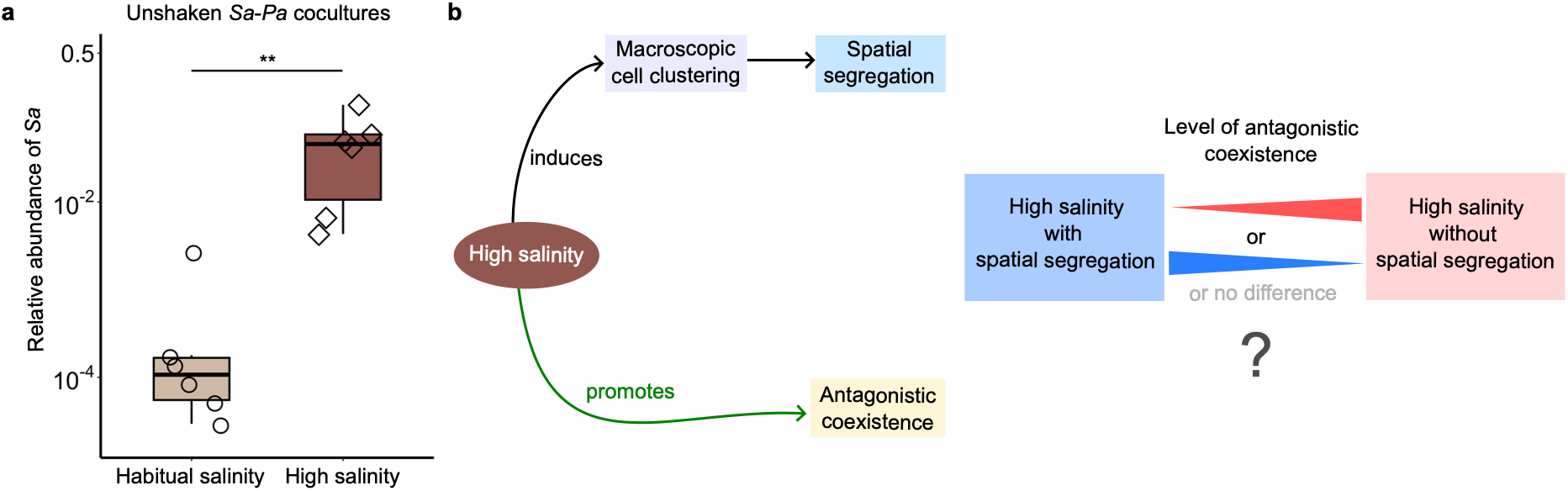
High salinity promotes the coexistence of *S. aureus* and *P. aeruginosa*. (a) The fraction of Sa after seven days of growth in two different environmental conditions (the inoculum *Sa*-*Pa* ratio was 1:1 for all microcosms). * denotes *P_adj_* < 0.05 (see *Statistical Appendix*). (b) A schematic of the key question arising from a combination of our observations in Figs. 1 and 2a: Does high salinity lead to better *Sa*-*Pa* coexistence *because* it spatially segregates the two species?

### Salinity-induced macroscopic clustering undermines *Sa-Pa* coexistence despite driving spatial segregation

To disentangle the role of the inducing environment from that of the induced phenotype in shaping *Sa-Pa* coexistence, we introduced a high salinity control that does not allow the spatial segregation of *Sa* and *Pa* (Fig. 3a). To this end, we vortexed the culture tubes for 20 s every six hours, which ensured that these control cultures were static for 99.9% of their incubation while also physically inhibiting *Sa*-*Pa* segregation (see *Methods*; Fig. S2). Such intermittent vortexing is an effective control here because it does not affect *Pa* or *Sa* cell counts on its own, as we show in detail later (also see *Methods*). We also included a similar control with intermittent vortexing under habitual salinity (Fig. 3a). Such intermittent vortexing, which allows cultures to remain static for 99.9% of the incubation, should lead to greater interference competition via active contact-based killing by increasing *Sa-Pa* intercellular contact and inhibiting spatial segregation (if any). Thus, if high salinity led to better *Sa-Pa* coexistence *because* it causes their spatial segregation, our high salinity control without spatial segregation should not exhibit such improved coexistence. Counterintuitively, we observed the highest level of *Sa-Pa* coexistence in high salinity *without* spatial segregation (Fig. 3b): here the *Sa-Pa* ratio after three days of coculture was almost identical to the inoculum *Sa-Pa* ratio (1:1). Moreover, high salinity without segregation led to a >100-fold higher median relative abundance of *Sa* than that in high salinity with segregation (and >1000-fold higher median relative abundance of *Sa* than that in habitual salinity; Fig. 3b). Hence, spatial segregation caused by macroscopic clustering cannot explain why high salinity leads to increased *Sa-Pa* coexistence (Fig. 3c). Furthermore, intermittent vortexing in habitual salinity led to decreased *Sa*-*Pa* coexistence, which aligns with the expectation of greater interference competition due to increased intercellular contacts (Fig. 3b). We observed a similar overall pattern when we repeated this series of coculture experiments with 7 days of incubation (Fig. S3). Taken together, although high salinity enhances *Sa-Pa* coexistence (Figs. 2 and 3b), it also induces phenotypically plastic macroscopic clustering that undermines such increase in coexistence (Figs. 3b and S3).

**Fig. 3.**
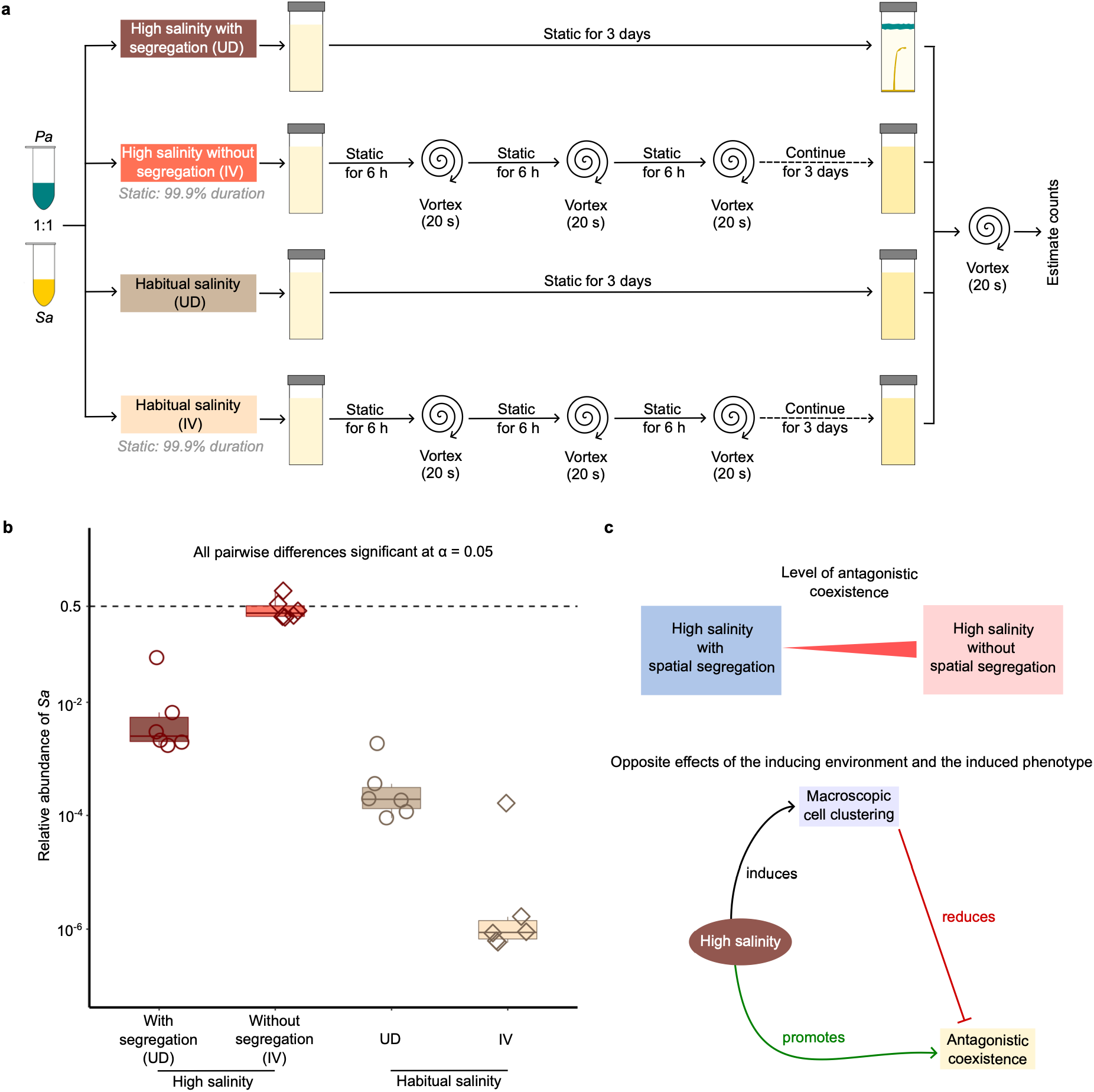
Disentangling the roles of the inducing environment and the induced phenotype in shaping *Sa*-*Pa* coexistence. (a) A schematic of our experimental workflow to determine if salinity-induced spatial segregation leads to higher *Sa*-*Pa* coexistence. UD and IV refer to ‘undisturbed’ and ‘intermittent vortexing’, respectively. (b) The level of coexistence after three days of growth under four distinct environmental conditions. High salinity without segregation showed the highest level of *Sa*-*Pa* coexistence. (*Sa* was undetectable in 5/6 replicates in Habitual salinity IV, see *Statistical Appendix* for the basis of our estimates of *Sa* counts in these replicates). (c) A schematic summary of our key finding: Spatial segregation cannot explain why high salinity leads to higher levels of *Sa*-*Pa* coexistence because macroscopic cell clustering reduces the level of achievable coexistence in high salinity.

To probe into why high-salinity-induced macroscopic clustering diminishes *Sa*-*Pa* coexistence, we studied how inhibiting such clustering itself shapes the growth of *Pa* and *Sa* in *monocultures*.

### Phenotypically plastic macroscopic clustering is adaptive for *Pa*, but not for *Sa*

We grew distinct *Pa* and *Sa* monocultures for three days in both high and habitual salinity, with and without intermittent vortexing for 20 s every six hours (see *Methods*). We found that inhibiting macroscopic clustering in high salinity led to a significant decrease in the total cell counts for *Pa* (Fig. 4a), but not for *Sa* (Fig. 4b). Put differently, in terms of total cell counts, high salinity-induced macroscopic cell clustering was adaptive for *Pa* (Fig. 4a), but neutral for *Sa* (Fig. 4b). The substantial reduction in *Pa* counts caused by inhibiting mat formation at the air-liquid interface can be readily intuited by invoking a reduced access to oxygen, which we revisit in the *Discussion*. Importantly, intermittent vortexing on its own cannot explain the differences in cell counts in our monoculture experiments. This is because in habitual salinity environments (where no macroscopic clustering is induced), intermittent vortexing for 20 s every six hours did not lead to any significant difference in the total cell counts of either *Sa* (*P_adj_* = 0.156) or *Pa* (*P_adj_* = 0.589) in monocultures (see *Statistical Appendix*).

Taken together, the high salinity environment not only induced contrasting morphologies of macroscopic clusters across *Pa* and *Sa* (Figs. 1 and S1), but such induced clustering also had contrasting adaptive value for these bacteria (Fig. 4). Our observations from these monocultures experiments led us to make three distinct serial predictions for *Sa-Pa* cocultures: ***Prediction 1*:** Similar to our observations in *Pa* monocultures, physically inhibiting the formation of high-salinity-induced *Pa* mats (by intermittent vortexing) should lead to reduced *Pa* counts in *cocultures* grown in high salinity. ***Prediction 2:*** Such reduction of *Pa* counts should lead to an increase in *Sa* counts in high salinity cocultures in a negatively correlated manner. ***Prediction 3:*** Given the lack of environmentally induced cell clustering in habitual salinity, no such negative correlation would be found across *Sa* and *Pa* counts in habitual salinity cocultures.

**Fig. 4.**
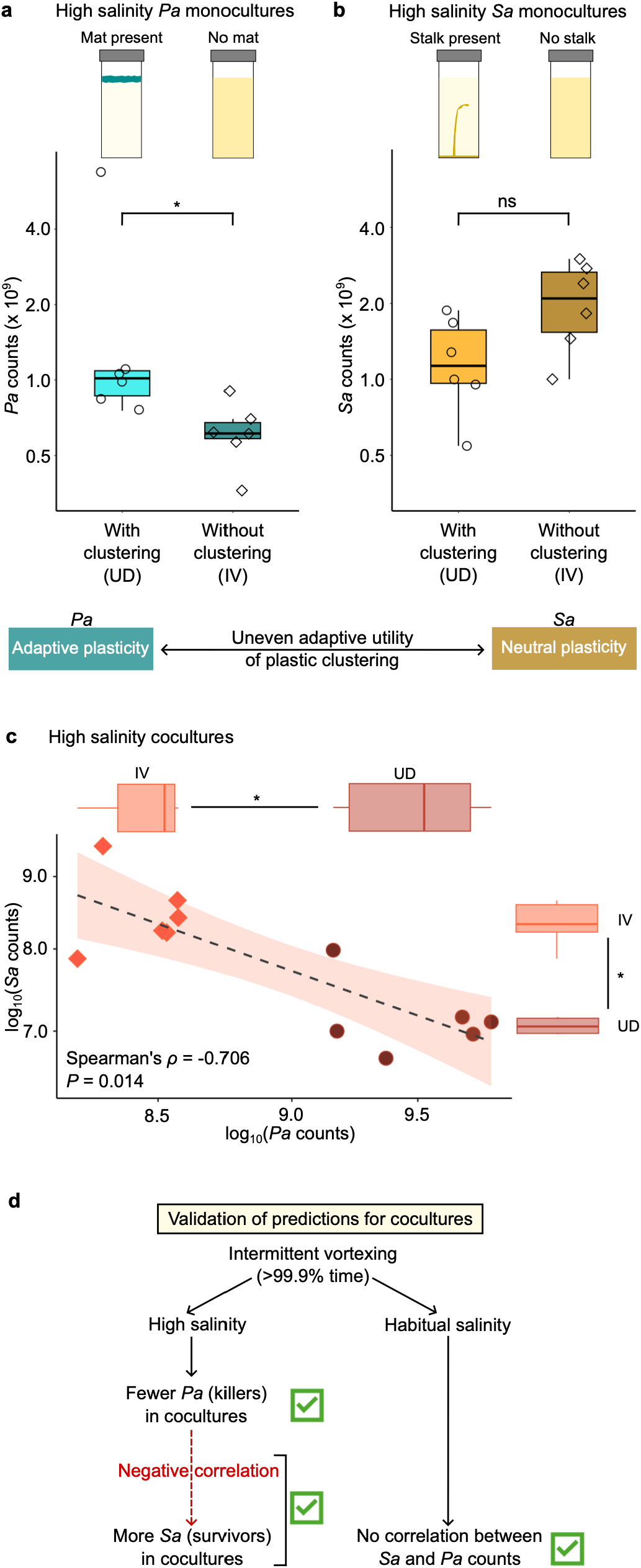
A quantitative explanation for why high salinity promotes coexistence when segregation is inhibited. (a) *Pa* cell counts after 3 days of growth in high salinity monocultures, with and without salinity-induced mat formation; UD and IV refer to ‘Undisturbed’ and ‘Intermittent vortexing’, respectively. * denotes *P_adj_* < 0.05 (see *Statistical Appendix* for details). (b) *Sa* cell counts after 3 days of growth in high salinity monocultures, with and without salinity-induced stalk formation; ns: not significant. In terms of total cell counts, plastic cell clustering is adaptive for *Pa*, but neutral for *Sa*. (c) A strong negative correlation between *Sa* and *Pa* counts in high salinity cocultures. (d) Our observations validated all of our serial predictions for high and habitual salinity cocultures. See Statistical *Appendix* for the details of the lack of significant correlation between *Sa* and *Pa* counts in habitual salinity cocultures.

To test the above serial predictions, we revisited our coculture experiments and analyzed them in terms of the total cell counts for *Pa* and *Sa* in both high and habitual salinity, with and without intermittent vortexing.

### Inhibiting plastic clustering leads to fewer killers and greater coexistence

Our experiments validated all of the above three serial predictions. Specifically, we found that in high salinity, intermittent vortexing (which inhibited spatial segregation) led to a substantial decrease in *Pa* counts and, simultaneously, to a significant increase in *Sa* counts (Fig. 4c). Moreover, as we had predicted, there was a strong negative correlation between *Sa* and *Pa* counts in high salinity cocultures (Fig. 4c). Furthermore, validating our prediction, we found no such correlation in habitual salinity cocultures (Fig. 4d). Taken together, our observations establish the inhibition of environmentally induced macroscopic cell clustering in the inducing environment as a quantitative driver of *Sa*-*Pa* coexistence: it reduces the number of killers (*Pa*) within high salinity microcosms, which results in more *Sa* survivors, and thus enhances *Sa*-*Pa* coexistence. Next, we sought to find out if such quantitative modulation of *Sa-Pa* coexistence can be explicitly explained by the extent of *Sa* killing by *Pa* in the inducing (high salinity) environment, with or without the induced phenotype (cell clustering). Fig. 3b suggests that contact-based killing of *Sa* by *Pa* plays a minor role in high salinity. This is because intermittent vortexing for 20 s every six hours, which should increase the probability of *Pa*-*Sa* intercellular contact in cultures that remain static for 99.9% of the incubation period, is expected to facilitate contact-based killing. However, as revealed by Fig. 3b, such intermittent vortexing led to a substantially *higher* fraction of *Sa* in high salinity. Hence, any physical facilitation of contact-based killing provided by the removal of spatial *Sa-Pa* segregation by intermittent vortexing was unsubstantial in the segregation-driving high salinity environment. Given such minor role played by contact-based killing in high salinity, we tested if the extent of diffusion-based *Sa* killing by *Pa* could explain why inhibiting phenotypically plastic clustering leads to greater *Sa-Pa* coexistence in high salinity.

### Diffusion-based killing captures why adaptive plasticity undermines *Sa*-*Pa* coexistence

We subjected a known number of viable *Sa* cells to cell-free spent media derived from *Pa* monocultures grown in high salinity, either with or without intermittent vortexing (which allowed or inhibited mat formation, respectively). As a control, we also subjected a similar number of viable *Sa* cells to cell-free spent media derived from *Sa* monocultures. Upon treating *Sa* cells with such spent media for 24 h, we determined the percentage survival (Fig. 5; see *Methods*). Whereas treatments with the spent media derived from high-salinity-grown *Pa* monocultures resulted in substantial reduction of viable *Sa* cells, no significant reduction in *Sa* counts was observed upon treatment with *Sa*-derived spent media control (Fig. 5). Crucially, we found that the high salinity-derived spent media form *Pa* killed *Sa* significantly more when the *Pa* monocultures were allowed to form high-salinity-induced mats as compared to when such mat formation was inhibited by intermittent vortexing (Fig. 5). This mechanistically corroborates the quantitative link (negative correlation) between *Pa* and *Sa* counts we had observed in high salinity *cocultures* (Fig. 4c). Taken together, when high-salinity-induced macroscopic mats are allowed to form, they lead to higher *Pa* counts and enable greater diffusion-based killing of *Sa,* thereby undermining *Sa-Pa* coexistence. Thus, a reduction in diffusion-based killing can explain why inhibiting the high-salinity-induced cell clustering in high salinity conditions (and abolishing *Sa-Pa* segregation as a consequence) leads to greater *Sa-Pa* coexistence.

**Fig. 5.**
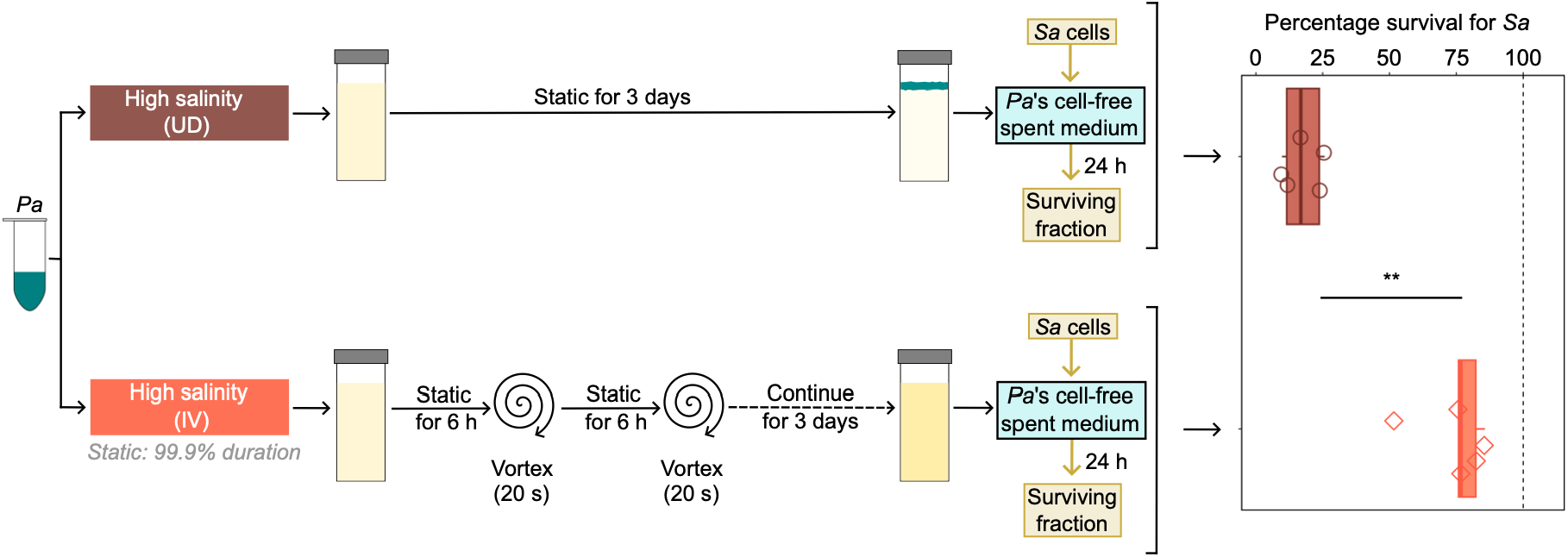
A mechanistic explanation for why high salinity promotes coexistence when segregation is inhibited. Spent media from high salinity *Pa* monocultures killed *Sa* significantly more when *Pa* mats were allowed to form. ** denotes *P* < 0.01 (see *Statistical Appendix*). *Sa* cells did not suffer any significant reduction in cell counts when they were subjected to spent media derived from *Sa* (*P* = 0.281 vs. 100% survival; see *Statistical Appendix*).

### Coexistence attained by inhibiting segregation enables rapid coculture-specific evolution

If our high-salinity-without-segregation treatment indeed provides ecologically significant increase in coexistence, it should also enable rapid molecular evolution comprising genomic changes selectively enriched in *Sa*-*Pa* cocultures, but not in monocultures. To test this prediction, we conducted pooled sequencing of cocultures grown for 7 days in high salinity without segregation and identified loci that showed mutations rising to a frequency of ≥ 10% (see *Methods*). We then determined if such loci also showed detectable mutations in *Sa* and *Pa* monocultures grown in high or habitual salinity, with or without intermittent vortexing for 7 days (16 pooled and 16 clonal sequences distributed over two species (*Sa* and *Pa*) in four distinct environmental treatments (Fig. 6a). Validating our prediction, two loci in *Pa* and one in *Sa* showed mutations consistent with coculture-specific evolution: mutations in these loci were selectively enriched to a frequency of ≥ 10% in the high-salinity-without-segregation cocultures, but were not detected in any of our 32 monoculture samples (Fig. 6a). Interestingly, the two loci in *Pa* that showed coculture-specific selective enrichment are known to shape key interactions with *S. aureus*^35,42–44^ (Fig. 6a). Thus, the stable *Sa*-*Pa* coexistence attained here provides a key biotic component of the environment (the presence of *Sa* cells in *Pa*’s habitat, and vice versa) that can shape rapid selective enrichment of *de novo* variation in a coculture-specific manner.

**Fig. 6.**
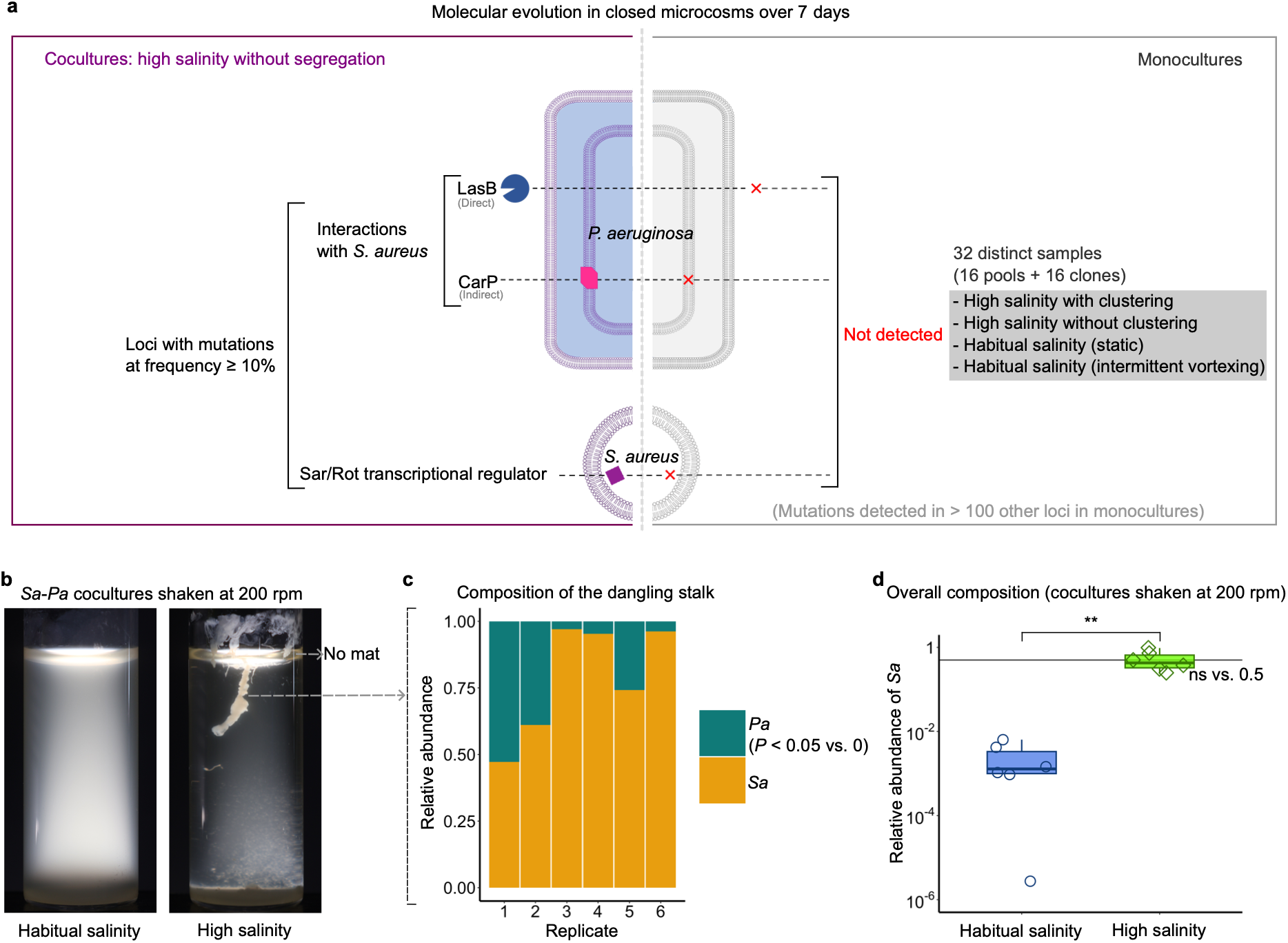
Validations of two key predictions emerging from our experiments. (a) Rapid coculture-specific molecular evolution enabled by inhibiting spatial segregation in high salinity. See *Sequencing Appendix* for the details of the detected mutations and *Methods* for the detectability criteria for mutations in pooled and clonal samples. (b) Rapid continuous shaking results inhibited mat formation in high salinity, but resulted in chimeric *Sa*-*Pa* clusters and substantially improved coexistence; ns: not significant; ** denotes *P* < 0.01 (see *Statistical Appendix* for details).

### Chimeric coexistence via the removal of oxygen monopoly

Having uncovered that inhibiting the formation of interface-inhabiting *Pa* mats via intermittent vortexing of static tubes improves *Sa-Pa* coexistence in high salinity, we hypothesized that another distinct physical means of inhibiting mat formation, namely, continuous shaking at high speed, would also lead to high levels of coexistence. Moreover, shaking coculture tubes continuously at high speed should not only prevent the formation of interface-inhabiting *Pa* mats, but it should also make oxygen more uniformly available than in static cultures, which should culminate in increased coexistence. To test this hypothesis, we initiated *Sa*:*Pa*∷1:1 cultures in both high and habitual salinity environments, and shook the culture tubes continuously at 200 rpm for 3 days (see *Methods*). As predicted, continuous shaking inhibited the formation of a horizontal mat covering the air-liquid interface in high salinity microcosms (Fig. 6b). Instead, each shaken high salinity culture exhibited a vertical stalk dangling from (but not entirely covering) the interface (Fig. 6b). Surprisingly, this stalk contained millions of *Sa* and *Pa* cells (Fig. 6c). Validating our predictions, the high salinity shaken cultures maintained 1:1 *Sa-Pa* ratios after three days of growth (Fig. 6d), but the habitual salinity shaken cultures (which were planktonic) failed to do so (Fig. 6d). The presence of a chimeric solid filamentous cluster underpins the unimportance of spatial segregation in improving *Sa-Pa* coexistence in high salinity. Our experiment also shows that removing *Pa*’s oxygen monopoly can be a key way of achieving high levels of coexistence.

## Discussion

Our study commences with the empirical demonstration that phenotypic plasticity can segregate antagonistic bacterial species in space (Fig. 1). Specifically, we show that a plastic phenotype (here, macroscopic cell clustering) induced by a common abiotic environmental cue (here, high salinity) can lead to the macroscopic segregation of two antagonistic bacterial species., *S. aureus* and *P. aeruginosa* (Fig. 1). This finding itself represents a key advance in understanding the significance of phenotypic plasticity: While plasticity is known to be an important factor in shaping ecological interactions^1,26,45,20,21^, empirical demonstrations of spatial segregation as a plasticity-driven phenomenon remain limited and indirect, particularly in the context of antagonistic species^26,27^. Moreover, while plasticity is known to influence multiple spatial aspects of microbial growth^46–50^, a clear elucidation of how plasticity causes the segregation of mutually antagonistic microbial species in space has remained elusive. Our study overcomes this key gap by showing how, by inducing macroscopic cell clustering, high salinity spatially segregates *Sa* and *Pa* (Fig. 1).

Ecological intuition suggests that such macroscopic segregation of *Sa* and *Pa* should weaken their interactions and promote their coexistence. Indeed, extensive theoretical^37,51–53^ and empirical^38,39,54,55^ studies have examined how such segregation reduces antagonistic interspecies interactions by limiting spatial overlap, thereby promoting coexistence. Moreover, given the generic applicability of spatial segregation in the context of both macro-^22,38,54^ and microorganisms^39,55^, it serves as a major ecological explanation for antagonistic coexistence^38^. Such segregation can arise from diverse ecological phenomena, including (but not limited to) constrained dispersal^56–58^, cooperation^59^, contact-mediated interference^60–63^, and inducible cell clustering (as revealed by our study). Crucially, these ecological drivers can link spatial reorganization to changes in growth, survival, and competitive interactions. Whether such linked effects heighten or undermine the overall antagonistic coexistence, beyond contributing to spatial patterning, is largely unknown. Importantly, whether an ecological driver of spatial segregation can itself undermine coexistence has remained unexplored. We address this key gap and show that ecological intuition can be highly misleading when it comes to spatial segregation and coexistence of antagonistic bacterial species.

Specifically, we tested whether the plasticity-driven *Sa-Pa* segregation *causes* an increase in the coexistence of these antagonistic bacteria (Fig. 3). A key strength of our study is the presence of an experimental control with the plasticity-inducing environment (high salinity) but without the induced phenotype (cell clustering). This control was implemented by physically inhibiting macroscopic clustering by intermittent vortexing of static tubes for 20 s every six hours. Importantly, such intermittent vortexing had no significant effect on either *Sa* or *Pa* counts on its own: specifically, in habitual salinity (an environment that does not induce cell clustering), intermittent vortexing did not significantly affect the monoculture counts of either *Sa* (*P_adj_* = 0.156) or *Pa* (*P_adj_* = 0.589; see *Statistical Appendix*). This control enabled us to test whether the induced phenotype is required for an increase in *Sa-Pa* coexistence or, parsimoniously, whether the inducing environment is sufficient *on its own* (*i.e.*, without the induced phenotype) for such increase. We found that high salinity could increase *Sa-Pa* coexistence on its own, and counterintuitively, high-salinity-induced macroscopic clustering undermines *Sa-Pa* coexistence despite spatially segregating *Sa and Pa* (Fig. 3). By explicitly disentangling the effects of the inducing environment from that of the induced phenotype, we reveal that even straightforward explanations based on the co-occurrence of a new (plastic) phenotype and an ecologically important change can be fallacious. Despite its expansive scale, spatial segregation itself becomes unimportant in our system because it is driven by an induced (plastic) phenotype that diminishes antagonistic coexistence. Such explicit causality-testing designs are rare in studies of phenotypic plasticity^3,4^, and are particularly difficult to perform in the context of interspecies coexistence^11,64^. A recent study *circumvented* this problem using ingenious experimental–theoretical approach and showed that plasticity can promote interspecies coexistence in plants^20^. Similarly, another recent study on interspecific coexistence in plants used a novel design to quantitatively disentangle direct competition from plastic interaction modification^21^. Employing a distinct yet powerful experimental design, here we reveal that plasticity can appear to promote antagonistic coexistence despite causally weakening it.

Using a series of experiments, we could successfully uncover the mechanistic basis of the opposing effects of the inducing environment and the induced phenotype on *Sa-Pa* coexistence (Figs. 4-6). We found that phenotypically plastic macroscopic clustering was adaptive for *Pa*, but not for *Sa* (Fig. 4). Previous studies with *Pseudomonas fluorescens* have shown that the formation of a thick mat at the air-liquid interface leads to an improved access to oxygen that provides a growth advantage under static conditions^65–67^. This can readily explain why, in our study, *Pa* achieved lower cell counts when the high-salinity-induced mat formation was inhibited by intermittent vortexing in monocultures that remained static for 99.9% of their incubation period (Fig. 4a). Aligning with this premise, we could achieve a similar modulation of *Pa* counts by either allowing or inhibiting mat formation under high salinity in *Sa-Pa* cocultures, which also ended up modulating the *Sa* counts in a negative correlated manner (Fig. 4c). Strengthening our explanation, no such correlation was found in the uninduced (habitual salinity) environment where no cell clustering occurs (Fig. 4d). Furthermore, consistent with our data on *Pa* cell counts, we found that in high salinity, *Pa*’s cell-free spent medium kills *Sa* more if the phenotypically plastic mats are allowed to form (Fig. 5). This provides a mechanistic backing to our quantitative explanation for why physically inhibiting high-salinity-induced clustering (which removes spatial segregation) improves *Sa*-*Pa* coexistence. Indeed *Pa* is known to diminish *Sa* in coculture using diverse diffusion-based interference mechanisms^31^, including phenazines^68^, 2-heptyl-4-quinolone N-oxide (HQNO)^33^, LasA protease^69^, inorganic polyphosphate^35^, rhamnolipids^70^, etc. Such diversity of diffusion-based mechanisms also supports the unimportance of spatial segregation in shaping *Sa*-*Pa* coexistence that we report here. Aligning with this broad notion, a recent study found that a lack of spatial structure can lead to improved interspecies coexistence in experimental bacterial communities^71^, and a diffusion-based mechanism was instrumental in that study’s context.

Our results also point towards the possibility that occupying the air-liquid interface can allow *Pa* to indirectly suppress *Sa*’s growth by monopolizing oxygen availability. Specifically, we removed any potential oxygen monopoly of *Pa* by shaking *Sa-Pa* cocultures continuously at 200 rpm: this not only ensured better aeration within the culture tube, but also inhibited the formation of a horizontal interface-covering *Pa* mat (Fig. 6b). Such continuous shaking resulted in the formation of a vertical macroscopic *Sa-Pa* chimera and led to a >100-fold increase in the median *Sa-Pa* coexistence relative to static high salinity conditions that allow *Sa-Pa* segregation (compare Fig. 6d with Fig. 3b). Thus, inhibiting the formation of *Pa* mats, either via intermittent vortexing (Fig. 3) or via continuous shaking (Fig. 6) not only prevents *Pa* from monopolizing oxygen by covering the air-liquid interface, it also promotes *Sa*-*Pa* coexistence.

*S. aureus* and *P. aeruginosa* are opportunistic pathogens that routinely co-infect the human respiratory tract^30–32^. Such coexistence in their natural habitats notwithstanding, *Sa* and *Pa* are extremely difficult to coculture in standard laboratory environments because *Pa* actively kills *Sa* and drives it to extinction^31–35^. Moreover, multiple studies have shown that some specific environmental conditions have the potential to improve the degree of *Sa*-*Pa* coexistence^31,72,73^. Hence, this species pair is an ideal system for studying how environmentally induced plastic phenotypes can shape the coexistence of mutually antagonistic species. Moreover, our study focuses on a plastic phenotype that provides a spatially explicit dimension to the ecological interactions between a pair of antagonistic species (where one species (*Pa*) can actively swim, but the other cannot). These aspects make our study system an ideal platform to test inferences about the precise role of plasticity-driven spatial structure in simple communities.

Taken together, our experiment comprehensively shows how plasticity can appear to increase antagonistic coexistence despite causally undermining it because an altered environment can increase coexistence independent of the plastic phenotype it induces. Thus, our experimental results fundamentally enhance the current understanding of the nuanced ecological roles phenotypic plasticity can play, even in the simplest of ecological communities.

## Methods

### Bacterial strains and culture conditions

We used the Gram-negative bacterium *Pseudomonas aeruginosa* (strain PAO1) and the Gram-positive bacterium *Staphylococcus aureus* (strain NCIM 5345 (ATCC 6538)) in our experiments. We cultured these bacteria in liquid environments within sealed glass tubes at 37°C (culture volume: 5 ml), either in monoculture or coculture (see below). We used two distinct kinds of nutrient media: (1) *Habitual salinity:* Luria Bertani (LB) broth, Miller (10 g/L tryptone, 5 g/L yeast extract, 10 g/L NaCl), (2) *High salinity:* Modified LB broth (10 g/L tryptone, 5 g/L yeast extract, 60 g/L NaCl). Depending on the growth treatment under consideration, the cultures were kept static, intermittently vortexed (for 20 s every six hours), or shaken continuously (see below for the details of specific growth treatments).

### Coculture and monoculture experiments in static conditions

We grew distinct clonal liquid monocultures derived from a single colony each of *Pa* and *Sa* in LB broth shaken at 200 rpm for 18 hours, estimated the number of bacteria in each monoculture via quantitative plating, and subsequently prepared several 1 ml cryo-stocks from each culture. Next, we initiated all coculture experiments in both habitual and high salinity with 1:1 ratio of *Sa* and *Pa*, with each bacterium starting with ∼5.4 million cells derived from freshly thawed cryo-stocks. Similarly, all the monoculture experiments were initiated with an identical number of *Sa* or *Pa*. Cryostocks once thawed were not used again to initiate a similar experiment. The culture tubes were incubated for either 3 days (Figs. 1 and 2) or 7 days (Fig. 3), after which the composition of each tube was estimated by quantitative plating (see below). We used Canon R50 (Canon Inc., Ōta, Japan) to capture backlit macroscopic images of culture tubes.

### Estimation of coculture composition

We determined the overall composition of cocultures by vortexing the culture tubes at 2800 rpm (orbital diameter: 4 mm) and subsequently determining the total colony forming units (CFUs) belonging to *Sa* and *Pa* by plating serial dilutions on two distinct media: (1) Luria Bertani agar (2) Staphylococcus agar no. 110 w/ azide. The latter does not allow the growth of *Pa*, but allows *Sa* to grow without inhibition. This dual plating approach enabled us to detect *Sa* in cocultures even if its abundance was a million-fold lower than that of *Pa*. For monocultures, such quantitative plating was done only on LB agar.

In cocultures, we used the relative abundance of *Sa*, which equals *N_Sa_*/(*N_Sa_* + *N_Pa_*), as a measure of coexistence. This usage is motivated by previous studies in microbial ecology^40,41^. Such a continuous relative-abundance-based measure is particularly relevant in quantifying coexistence when multiple experimental treatments are expected to show highly asymmetric abundances across the two species. This measure is also functionally analogous to the Gini coefficient for economic inequality across two categories, and can be used similarly for determining the evenness of the distribution between the two species.

### Estimation of mat and stalk composition in cocultures

From a given three-day-old *Sa*-*Pa* high salinity coculture, we first extracted the mat by spooling it using a plastic inoculation loop. Next, we isolated a small piece of the spooled mat fitting within 20 µl using a micropipette and delivered it into a total of 2 ml fresh medium in a 24 well plate. This clustered piece of the mat was then subjected to four serial washes in 2 ml fresh media in four distinct wells, which resulted in at least 10^-8^-fold dilution of any planktonic bacteria while keeping the clustered piece of the mat undiluted. Next, we added the serially washed mat cluster (fitting within 20 µl) to 180 µl of LB, vortexed it vigorously and immediately subjected it to quantitative dual-media plating (as described above) to estimate the mat’s composition. We used a similar approach to subsequently extract and wash the stalk serially before subjecting it to quantitative dual-media plating to determine its composition.

### Physical inhibition of macroscopic clustering and spatial segregation

We initiated 1:1 *Sa*-*Pa* cocultures along with the two monoculture controls in both high and habitual salinity and grew them for 3 days, as described above. In both the environments, one set of cultures remained static throughout their incubation while another set underwent vortexing for 20 seconds every six hours (Fig. 3). Such intermittent vortexing ensured that these cultures were static for 99.9% of their incubation while also physically inhibiting *Sa*-*Pa* segregation (which occurs only in high salinity). Such physical inhibition of clustering (and thus *Sa*-*Pa* segregation) enabled us to disentangle the effects of the inducing environment (high salinity) from that of the induced phenotype (macroscopic clustering). Such physical inhibition is effective in our experiments because it does not lead to any significant change in *Pa* or *Sa* cell counts on its own. Specifically, in habitual salinity (an environment that does not induce cell clustering), intermittent vortexing does not change the cell counts in either *Sa* (*P_adj_* = 0.156) or *Pa* (*P_adj_* = 0.589) monocultures, nor does it change the total cell counts in *Sa*-*Pa* cocultures in habitual salinity (*P* = 0.937). Thus, such physical intervention, which does not affect *Sa* or *Pa* counts directly, allows us to incorporate a control that offers the inducing environment without the induced phenotype.

### Diffusion-based killing assays

We grew *Pa* monocultures in high salinity, either with or without intermittent vortexing (referred to as IV and UD, respectively) for three days (8 replicate glass tubes for each treatment; culture volume = 5 ml). Subsequently, for each treatment (UD and IV), we pooled the 8 replicate cultures and centrifuged the pooled contents for 10 minutes at 5000 rcf at 4°C, retained the supernatant, and discarded the pellet. The retained supernatant was subjected to a similar round of centrifugation. The supernatant obtained after the second centrifugation was then subjected to two successive rounds of filter sterilization using filters with pore size 0.2 µm to obtain cell-free spent media from *Pa*. We plated undiluted spent media on LB agar plates to verify that it was devoid of any viable *Pa* cells; if a *Pa* colony was detected on LB agar, the spent medium was subjected to another round of filter sterilization. Next, the cellfree spent medium for each treatment was redistributed randomly into 5 wells within a 96 well plate. We subjected a predetermined number of freshly grown *Sa* cells (derived from a single colony and grown for 24 hours in habitual salinity under shaken conditions) to the above two *Pa*-cell-free spent medium treatments (UD and IV) for 24 hours. As an additional control, we also subjected the freshly grown *Sa* cells to spent media derived from *Sa*. In each case, we determined *Sa* counts after 24 h to obtain the percentage survival of *Sa*.

### Inhibiting *Pa* mat formation by the continuous shaking of cocultures

We initiated 1:1 *Sa*-*Pa* cocultures and subjected them to continuous orbital shaking at 200 rpm (orbital diameter: 26 mm) for 3 days. This inhibited the formation of a horizontal mat covering the air-liquid interface, but allowed the formation of a vertical stalk dangling from the interface (Fig. 6b). To determine the composition of this stalk, we used a similar approach to that used for determining the composition of mats and stalks in static cocultures described above.

### Statistics

We used non-parametric statistics throughout our analyses. The details of the individual statistical analyses for each experiment are given in *Statistical Appendix*.

### Community-wide genomics

We subjected the following samples to whole genome sequencing: Ancestral *Sa* (clone) and *Pa* (clone), 7-day old high salinity cocultures without spatial segregation (pools), *Sa* and *Pa* monocultures grown in high or habitual salinity, with or without intermittent vortexing for 7 days (both clones and pools; see *Sequencing Appendix* for details). We outsourced the extraction of DNA, library preparation, and sequencing to an external vendor (Nucleome Informatics Pvt. Ltd., India). The genomic DNA was isolated from each sample using the DNeasy PowerSoil Pro kit (QIAGEN Inc., USA), the quantity of extracted DNA was determined using Qubit 4 (Thermo Fisher Scientific, USA), and the DNA’s purity was verified using A260/A280 and A260/A230 ratios using Nanodrop 2000 (Thermo Fisher Scientific, USA). KAPA HyperPlus Kit (Roche Molecular Systems, Inc., USA) was used to prepare DNA libraries, and Agilent 2100 Bioanalyzer (Agilent Technologies Inc., USA) was used for the library length assessment. 250 bp paired-end next generation sequencing was performed with the prepared libraries using Illumina NovaSeq 6000 (Illumina Inc., USA). We used Geneious Prime (GraphPad Software LLC, USA) to analyze the sequencing output and trimmed the raw sequence data using BBDuk to remove reads < 20 bp or with a quality score < 20. To identify mutations, we compared the evolved samples to their ancestral sequences, and restricted our analysis to variants with the following features: coverage: ≥ 100x for pools and ≥ 20x for clones; variant frequency: ≥ 10% for pools and ≥ 80% for clones; strand bias ≤ 70%; sequencing quality ≥ 30. We also manually inspected the mutated contigs qualifying these criteria and checked for mapping anomalies. We did not include intergenic regions and synonymous mutations in our analyses. We assessed if the high level of coexistence (*Sa*:*Pa*∷∼1:1) enabled by inhibiting segregation in high salinity cocultures also allowed coculture-specific evolution. To this end, we determined if the loci with mutations rising to ≥ 10% in these cocultures were also found to have detectable mutations in our monoculture samples in accordance with the above criteria for pools and clones (Fig. 6a).

## Supporting information

Statistical Appendix

Sequencing Appendix

## Acknowledgements

We thank Will Ratcliff, Sutirth Dey, Ullasa Kodandaramaiah, and Hema Somanathan for valuable discussions and constructive critiques. We are grateful to Shraddha Karve and Sunish Kumar Radhakrishnan for their help with bacterial strains. This work was supported by intramural funds provided by IISER Thiruvananthapuram to Y.C., who also acknowledges extramural support from ANRF (grant ANRF/ECRG/2024/001946/LS). D.E.S. was supported by an INSPIRE undergraduate fellowship, sponsored by Department of Science and Technology (DST), Govt. of India. M.H. was supported by a Senior Research Fellowship sponsored by Council for Scientific and Industrial Research (CSIR), Govt. of India. The funders had no role in study design, data collection and analysis, preparation of the manuscript, or the decision to publish.

## AI disclosure

We used ChatGPT and Gemini for language refinement and incorporated minor linguistic suggestions made by AI after thorough review. No data was generated using AI. We take full responsibility for all the findings reported here.

## Author contributions

Conceived the original idea and designed the project: Y.C.

Conducted the experiments and data analysis: D.E.S. and M.H.

Supervised the project: Y.C.

Wrote the manuscript: Y.C. and M.H.

Acquired funding: Y.C.

## Data Availability

All the data reported here are included in the manuscript and its Supplementary Information files, except for the raw whole-genome sequencing data files, which will made available publicly upon publication of the peer-reviewed article.

## Competing interests

The authors declare no competing interests.

## Notes

### Competing Interest Statement

The authors have declared no competing interest.

