## Supplementary material for "Phenotypic plasticity segregates antagonistic species but undermines their coexistence": Statistical Appendix

***to***

We used non-parametric statistics throughout our analyses. All the analyses were done with R (2022.02.3 Build 492). Standard R functions and scripts were used in the analyses.

**Contents**


| Section | Details |
| --- | --- |
| 1 | Statistical details for Fig. 1b |
| 2 | Statistical details for Fig. 2a |
| 3 | Statistical details for Fig. 3b |
| 4 | Statistical details for Fig. S3 |
| 5 | Statistical details for Fig. 4a |
| 6 | Statistical details for Fig. 4b |
| 7 | Statistical details for Figs. 4c and 4d |
| 8 | Statistical details for Fig. 5 |
| 9 | Statistical details for Figs. 6c and 6d |

**1: Statistical details for Fig. 1b**

***1.1***
Mat composition:

Percentage of *Pa* in the mats:
One-sided one-sample Wilcoxon signed-rank tests against the reference value of 100% (N = 6): $P = 0.091$.

***1.2***
Stalk composition:

Percentage of *Sa* in the stalks:
One-sided one-sample Wilcoxon signed-rank tests against the reference value of 100% (N = 6): $P = 0.091$.

**2: Statistical details for Fig. 2a**

Relative abundance of *Sa*:

Compared groups (N = 6 for each):

- Habitual salinity coculture
- High salinity coculture

Two-tailed Mann-Whitney U test: $P = 0.0021$

**3: Statistical details for Fig. 3b**

Compared groups (N = 6 for each):

- High salinity with segregation (UD)
- High salinity without segregation (IV)
- Habitual salinity (UD)
- Habitual salinity (IV)

Kruskal-Wallis-test: Omnibus $P = 0.0001104$.

Post-hoc pairwise comparisons: Two-tailed Mann-Whitney U tests followed by Holm-Bonferroni correction:

| Group pair | Raw *P* | Adjusted *P* (Holm-Bonferroni correction) |
|---|---|---|
| High salinity UD vs. high salinity IV | 0.002165 | 0.0129900000 |
| High salinity UD vs. habitual salinity UD | 0.004329 | 0.0129900000 |
| High salinity UD vs. habitual salinity IV | 0.002165 | 0.0129900000 |
| High salinity IV vs. habitual salinity IV | 0.002165 | 0.0129900000 |
| High salinity IV vs. habitual salinity UD | 0.002165 | 0.0129900000 |
| Habitual salinity UD vs. habitual salinity IV | 0.008658 | 0.0129900000 |

*Sa* was undetectable in 5/6 replicates in the habitual salinity (IV) treatment. Specifically, plating 100 µl of a $10^{-2}$ dilution (the most lenient dilution used in this assay) did lead to any detectable *Sa* colonies on solid plates with Staphylococcus agar no. 110 w/ azide for this treatment. However, since undiluted cultures were not plated due to logistical constraints, assigning zero *Sa* counts to this treatment would be misleading. Hence, we used the most conservative approach to assign *Sa* counts for these replicates for plotting and statistics: our approach minimizes the between-group differences. Specifically, since the habitual salinity (IV) treatment showed the lowest abundance of *Sa* among the four treatments, we assigned the highest theoretical count that will fail to get detected in quantitative plating of 100 µl of $10^{-2}$-diluted culture. Following this criterion, we assigned 99 *Sa* cells per 100 µl of $10^{-2}$-diluted habitual salinity (IV) treatment in replicates where *Sa* could not be physically detected. Despite this most conservative approach, all pairwise group differences were found to be significant ($P_{adj} < 0.05$; see above).

**4: Statistical details for Fig. S3**

Compared groups (N = 6 for each):

- High salinity with segregation (UD)
- High salinity without segregation (IV)
- Habitual salinity (UD)
- Habitual salinity (IV)

Kruskal-Wallis-test: Omnibus *P* = 0.00030.

Post-hoc pairwise comparisons: Two-tailed Mann-Whitney U tests followed by Holm-Bonferroni correction:

| Group pair | Raw *P* | Adjusted *P* (Holm-Bonferroni correction) |
|---|---:|---:|
| High salinity UD vs. high salinity IV | 0.002165 | 0.0129900000 |
| High salinity UD vs. habitual salinity UD | 0.002165 | 0.0129900000 |
| High salinity UD vs. habitual salinity IV | 0.01515 | 0.0303000000 |
| High salinity IV vs. habitual salinity IV | 0.002165 | 0.0129900000 |
| High salinity IV vs. habitual salinity UD | 0.002165 | 0.0129900000 |
| Habitual salinity UD vs. habitual salinity IV | 0.3939 | 0.3939 |

Since *Sa* was undetectable in 2/6 replicates in Habitual salinity (IV) treatment, we used the same approach as that for Fig. 3 (see section 3 above) to most conservatively estimate and assign *Sa* counts for these replicates.

**5: Statistical details for Fig. 4a**

*Pa* counts in monocultures

Compared groups (N = 6 for each):

- High salinity with segregation (UD)
- High salinity without segregation (IV)
- Habitual salinity (UD)
- Habitual salinity (IV)

Kruskal-Wallis-test: Omnibus $P = 0.000368$.

Post-hoc pairwise comparisons: Two-tailed Mann-Whitney U tests followed by Holm-Bonferroni correction:

| Group pair | Raw *P* | Adjusted *P* (Holm-Bonferroni correction) |
|---|---:|---:|
| High salinity UD vs. high salinity IV | 0.01291 | 0.025974 |
| High salinity UD vs. habitual salinity UD | 0.004329 | 0.025974 |
| High salinity UD vs. habitual salinity IV | 0.004329 | 0.025974 |
| High salinity IV vs. habitual salinity UD | 0.004998 | 0.025974 |
| High salinity IV vs. habitual salinity IV | 0.004998 | 0.025974 |
| Habitual salinity UD vs. habitual salinity IV | 0.5887 | 0.5887 |

**6: Statistical details for Fig. 4b**

*Sa* counts in monocultures

Compared groups (N = 6 for each):

- High salinity with segregation (UD)
- High salinity without segregation (IV)
- Habitual salinity (UD)
- Habitual salinity (IV)

Kruskal-Wallis-test: Omnibus $P = 0.0260$.

Post-hoc pairwise comparisons: Two-tailed Mann-Whitney U tests followed by Holm-Bonferroni correction:

| Group pair | Raw *P* | Adjusted *P* (Holm-Bonferroni correction) |
|---|---:|---:|
| High salinity UD vs. high salinity IV | 0.06494 | 0.3247 |
| High salinity UD vs. habitual salinity UD | 0.06494 | 0.3247 |
| High salinity UD vs. habitual salinity IV | 0.8182 | 0.8182 |
| High salinity IV vs. habitual salinity UD | 0.06494 | 0.3247 |
| High salinity IV vs. habitual salinity IV | 0.09307 | 0.3247 |
| Habitual salinity UD vs. habitual salinity IV | 0.02597 | 0.15582 |

**7: Statistical details for Figs. 4c and 4d**

***7.1***

Correlation between *Sa* and *Pa* counts in high salinity cocultures:
Spearman's correlation coefficient ($\rho$) = -0.706, $P$ = 0.014.

***7.2***

*Pa* counts in cocultures
Compared groups (N = 6 for each):

- High salinity with segregation (UD)
- High salinity without segregation (IV)
- Habitual salinity (UD)
- Habitual salinity (IV)

Kruskal-Wallis-test: Omnibus $P$ = 0.0028.

Post-hoc pairwise comparisons: Two-tailed Mann-Whitney U tests followed by Holm-Bonferroni correction:

| Group pair | Raw *P* | Adjusted *P* (Holm-Bonferroni correction) |
|---|---:|---:|
| High salinity UD vs. high salinity IV | 0.002165 | 0.01299 |
| High salinity UD vs. habitual salinity UD | 0.2403 | 0.7209 |
| High salinity UD vs. habitual salinity IV | 0.3095 | 0.7209 |
| High salinity IV vs. habitual salinity UD | 0.002165 | 0.01299 |
| High salinity IV vs. habitual salinity IV | 0.002165 | 0.01299 |
| Habitual salinity UD vs. habitual salinity IV | 0.9372 | 0.9372 |

***7.3***

*Sa* counts in cocultures
Compared groups (N = 6 for each):

- High salinity with segregation (UD)
- High salinity without segregation (IV)
- Habitual salinity (UD)
- Habitual salinity (IV)

Kruskal-Wallis-test: Omnibus $P = 0.8 \times 10^{-4}$.

Post-hoc pairwise comparisons: Two-tailed Mann-Whitney U tests followed by Holm-Bonferroni correction:

| Group pair | Raw *P* | Adjusted *P* (Holm-Bonferroni correction) |
|---|---:|---:|
| High salinity UD vs. high salinity IV | 0.004329 | 0.021606 |
| High salinity UD vs. habitual salinity UD | 0.004998 | 0.021606 |
| High salinity UD vs. habitual salinity IV | 0.003665 | 0.021606 |
| High salinity IV vs. habitual salinity UD | 0.004998 | 0.021606 |
| High salinity IV vs. habitual salinity IV | 0.003665 | 0.021606 |
| Habitual salinity UD vs. habitual salinity IV | 0.003601 | 0.021606 |

Note: *Sa* was undetectable in 5/6 replicates in Habitual salinity (IV) treatment, and following the most conservative approach (as described in section SA.3 above)

***7.4***

Correlation between *Sa* and *Pa* counts in habitual salinity cocultures:
Spearman's correlation not significant ($P = 0.432$).

Note: As described in section 3 above, *Sa* was undetectable in 5/6 replicates in the habitual salinity (IV) treatment, and following the most conservative approach, we assigned the *highest* value between 0 and 99 to the count of *Sa* cells per 100 µl of $10^{-2}$-diluted habitual salinity (IV) treatment in replicates where *Sa* could not be physically detected. Such assignment for these replicates notwithstanding, given the rest of our data based on *Sa* and *Pa* counts, it is mathematically trivial to show that it is impossible for any combination of five values between 0 and 99 (which is where the real *Sa* counts would lie in these replicates where *Sa* was undetected) to lead to a significant correlation between *Sa* and *Pa* counts here.

**8: Statistical details for Fig. 5**

***8.1***

Percentage survival for *Sa*:

Compared groups (N = 5 for each):

- Spent medium from *Pa* monoculture in High salinity (UD)
- Spent medium from *Pa* monoculture in High salinity (IV)

Two-tailed Mann-Whitney U test: $P = 0.0079$.

***8.2***

*Sa* percentage survival in its own spent medium over 24 h:
Two-sided one-sample Wilcoxon signed-rank test against the reference value of 100% (N = 5): $P = 0.2807$

### 9: Statistical details for Figs. 6c and 6d

***9.1***
Percentage of *Pa* in chimeric clusters:

One-sided one-sample Wilcoxon signed-rank test against the reference value of 0% (N = 6): $P = 0.018$

***9.2***
Relative abundance of *Sa* in shaken cocultures:

Compared groups (N = 6 for each):

- Habitual salinity shaken cocultures
- High salinity shaken cocultures

Two-tailed Mann-Whitney U test: $P = 0.0022$.

***9.3***
Relative abundance of *Sa* in shaken cocultures:
Two-sided one-sample Wilcoxon signed-rank test against the reference value of 0.5 (N = 6): $P \approx 1$.
