## Supplementary Information for "Phenotypic plasticity segregates antagonistic species but undermines their coexistence"

*for*

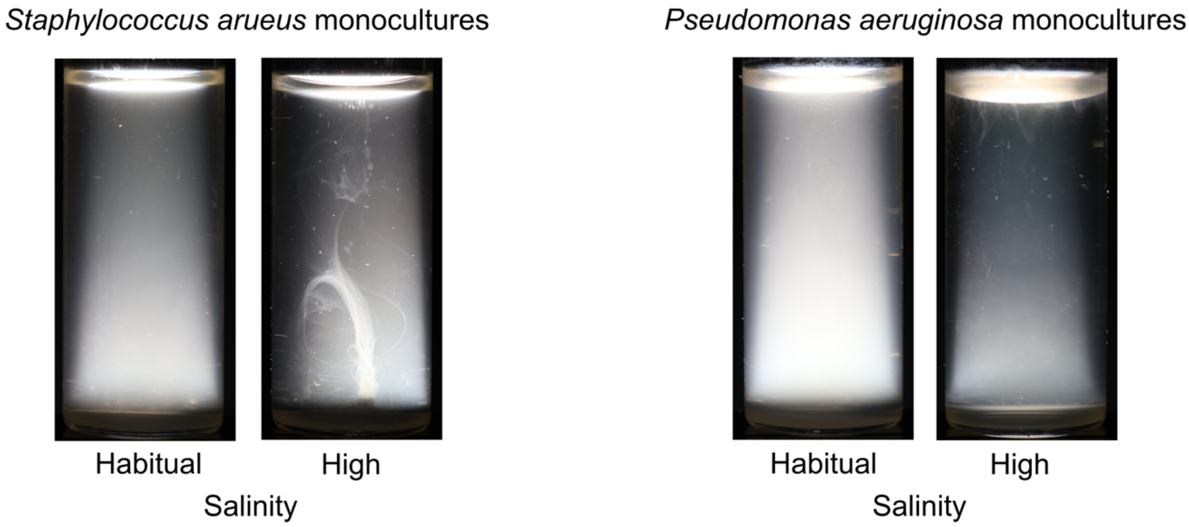


**Fig. S1. Phenotypically plastic macroscopic clustering of cells in *Staphylococcus aureus* (*Sa*) and *Pseudomonas aeruginosa* (*Pa*).** High salinity made *Sa* and *Pa* form markedly contrasting macroscopic clusters that differed in both morphology and spatial location.

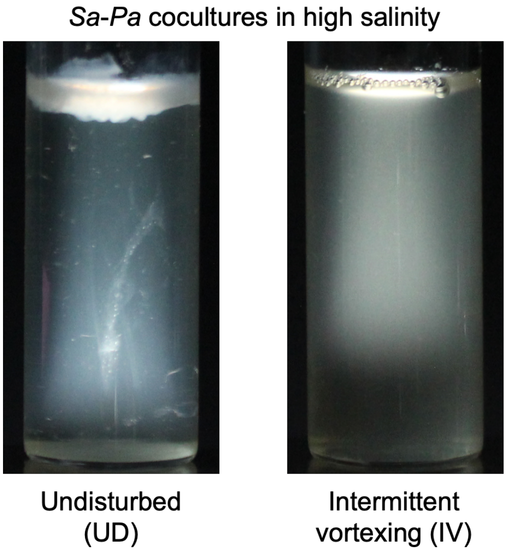


**Fig. S2. Intermittent vortexing inhibits cell clustering and abolishes spatial segregation in *Sa-Pa* cocultures.**

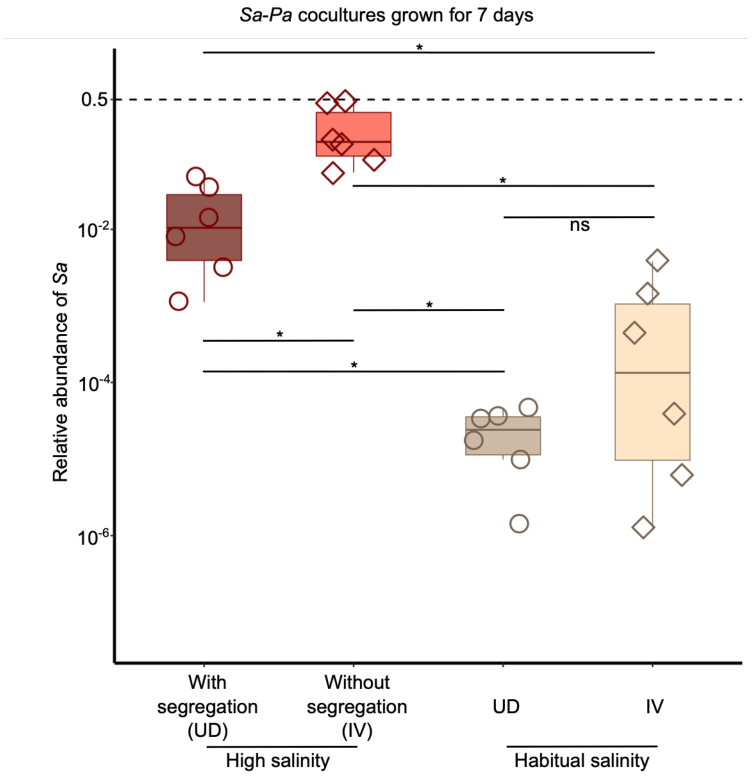


**Fig. S3. Disentangling the roles of the inducing environment and the induced phenotype in shaping *Sa-Pa* coexistence over 7 days.** High salinity without segregation showed the highest level of *Sa-Pa* coexistence. (*Sa* was undetectable in 2/6 replicates in Habitual salinity IV, see *Statistical Appendix* for the basis of our estimates of *Sa* counts in these replicates). * denotes $P_{adj} < 0.05$; ns: not significant (see in *Statistical Appendix*). In high salinity UD, from the fourth day onwards, all replicates showed an interface-dwelling greenish mat characteristic of *Pa*. The evolution of such mats in static cultures after four days is expected based on previous studies with *P. aeruginosa* because the mat-forming genotypes are adaptive in such environments.
